# The spindle protein CKAP2 regulates microtubule dynamics and ensures faithful chromosome segregation

**DOI:** 10.1101/2023.10.27.564280

**Authors:** Lia Mara Gomes Paim, Azriel Abraham Lopez-Jauregui, Thomas S. McAlear, Susanne Bechstedt

## Abstract

Regulation of microtubule dynamics by microtubule-associated proteins (MAPs) is essential for mitotic spindle assembly and chromosome segregation. Altered microtubule dynamics, particularly increased microtubule growth rates, were found to be a contributing factor for the development of chromosomal instability, which potentiates tumorigenesis. The MAP XMAP215/CKAP5 is the only known microtubule growth factor, and whether other MAPs regulate microtubule growth in cells is unclear. Our recent *in vitro* reconstitution experiments have demonstrated that Cytoskeleton-Associated Protein 2 (CKAP2) increases microtubule nucleation and growth rates, and here we find that CKAP2 is also an essential microtubule growth factor in cells. By applying CRISPR-Cas9 knock-in and knock-out as well as microtubule plus-end tracking live cell imaging, we show that CKAP2 is a mitotic spindle protein that ensures faithful chromosome segregation by regulating microtubule growth. Live cell imaging of endogenously-labelled CKAP2 showed that it localizes to the spindle during mitosis, and rapidly shifts its localization to the chromatin upon mitotic exit before being degraded. Cells lacking CKAP2 display reduced microtubule growth rates and an increased proportion of chromosome segregation errors and aneuploidy that may be a result of an accumulation of kinetochore-microtubule mis-attachments. Microtubule growth rates and chromosome segregation fidelity can be rescued upon CKAP2 expression in knock-out cells, revealing a direct link between CKAP2 expression and microtubule dynamics. Our results unveil a role of CKAP2 in regulating microtubule growth in cells and provide a mechanistic explanation for the oncogenic potential of CKAP2 misregulation.

**Significance statement:** Cell division is accomplished by the assembly of a mitotic spindle composed of microtubules that segregate the chromosomes. Cells with altered microtubule dynamics frequently mis-segregate chromosomes and develop aneuploidy, which contributes to cancer development. However, how microtubule dynamics are regulated in cells is not entirely understood. Here, using CRISPR-Cas9 genome editing and live cell imaging, we find that the microtubule-associated protein CKAP2 tightly regulates microtubule growth and ensures the proper segregation of chromosomes. Cells lacking CKAP2 develop errors in chromosome segregation and aneuploidy due to a substantial decline in microtubule growth rates. The essential role of CKAP2 in the regulation of microtubule growth provides an explanation for the oncogenic potential of CKAP2 misregulation.

**Classification:** Biological Sciences – Cell Biology

## Introduction

Mitotic cell divisions are accomplished when replicated chromosomes are successfully segregated into two new daughter cells. This process relies on the assembly of a bipolar spindle composed of microtubules that are nucleated by two centrosomes, followed by the attachment of microtubules to kinetochores and alignment of sister chromatids at the metaphase plate (1, 2). Upon chromosome alignment, anaphase is triggered, during which sister chromatids are pulled apart by spindle microtubules towards the cell poles (1). Mitosis is then completed by cytokinesis, when the cytoplasm is partitioned into two daughter cells that carry the segregated chromosomes (3). Accurate spindle assembly is key to faithful chromosome segregation. Altered microtubule dynamics can lead to chromosomal instability (CIN), where cells persistently exhibit chromosome segregation errors over multiple cell divisions resulting in aneuploidy, both of which contribute to tumorigenesis (4–6).

Throughout mitosis, microtubules nucleated from the centrosomes undergo continuous cycles of growth and shrinkage, a property known as dynamic instability (7) that is regulated by microtubule-associated proteins (MAPs) and motor proteins (8, 9). Microtubule growth rates have been found to be increased in chromosomally unstable cells and knockdown of the MAP chTOG/XMAP215/CKAP5 was shown to reduce microtubule growth rates in those cells and reduce CIN (5), although others have reported no impact of chTOG depletion upon microtubule growth in HeLa cells (10). Nevertheless, an increase in microtubule growth rates has been considered a stepping stone to CIN (5). On the other hand, microtubule growth rates were reduced in the presence of the MAP TPX2 in *in vitro* reconstitution assays (11), and overexpression of TPX2 in hTERT RPE-1 cells leads to spindle assembly and nuclear defects (12), perhaps suggesting that excessively low microtubule growth rates may also impact chromosome segregation fidelity. However, whether an excessive reduction in microtubule growth rates in cells can also cause chromosome mis-segregation and aneuploidy has not been formally explored. In addition, chTOG/XMAP215/CKAP5 is the only known microtubule polymerase (13), and whether other MAPs regulate microtubule growth in cells is unclear.

The Cytoskeleton-Associated Protein 2 (CKAP2) is a MAP expressed during mitosis that localizes to centrosomes and spindles before being targeted for degradation by the APC/C at mitotic exit (14, 15). Previous work has shown that CKAP2 is a spindle protein involved in ploidy maintenance and chromosome segregation (16–19). CKAP2 knock-down leads to spindle assembly defects (16) and CKAP2 overexpression has been associated with tumor development (20–22) and is negatively correlated with survival rates in cancer patients (23). Our recent *in vitro* reconstitution experiments have demonstrated that CKAP2 substantially increases microtubule nucleation and growth rates (24), however, whether and how CKAP2 regulates microtubule dynamics and chromosome segregation in cells has yet to be explored. Here we take advantage of CRISPR-Cas9 knock-in and knock-out approaches to address the roles of CKAP2 in cells. Using extensive live cell imaging and plus-end microtubule tracking experiments, we find that CKAP2 is a microtubule growth factor in cells that is essential for regulating microtubule dynamics during mitosis and ensuring faithful chromosome segregation.

## Results

### Cell cycle-dependent expression and subcellular localization of CKAP2

Previous immunofluorescence studies have demonstrated that CKAP2 localizes primarily to the spindle during mitosis (25, 26). To further investigate the temporal dynamics of CKAP2 expression and localization, we performed genome editing using CRISPR-Cas9 knock-in to introduce an N-terminal GFP-tag to CKAP2 in hTERT-immortalized retinal pigment epithelial cells (RPE-1) and HT1080 cells, and assessed CKAP2 expression and localization with live cell and immunofluorescence imaging (Supplementary Figure 1a; Supplementary Video 1). We found that CKAP2 localizes to duplicated centrosomes and microtubules at late interphase (Fig. 1e-f) and early mitosis (nuclear envelope breakdown – NEBD), and it reaches its peak expression at metaphase, during which it localizes to the mitotic spindle (Fig. 1a and b; Supplementary Figure 1b; Supplementary Video 1). Exposure of GFP:CKAP2 metaphase cells to ice-cold treatment - which allows for the visualization of kinetochore fibers (k-fibers) - revealed that CKAP2 retains its microtubule localization after cold shock, remaining co-localized with k-fibers (Fig. 1g-h). Following anaphase onset, CKAP2 undergoes a gradual shift in localization, at first localizing to both microtubules and chromatin within 3.54 ± 1.18 min minutes of anaphase onset (Fig.1a mid-anaphase; 1c-d; Supplementary Video 1), followed by a complete displacement to chromatin within 20.45 ± 2.56 minutes of anaphase onset (Fig.1a telophase; 1c-d; Supplementary Figure 1b) before fluorescence values decrease to near-background levels (Fig. 1a-b; Supplementary Figure 1b; Supplementary Video 1). A similar pattern of CKAP2 expression and localization was observed with immunofluorescence (Supplementary Figure 1c-d) and is consistent with previous reports (25, 26).

**Figure 1.**
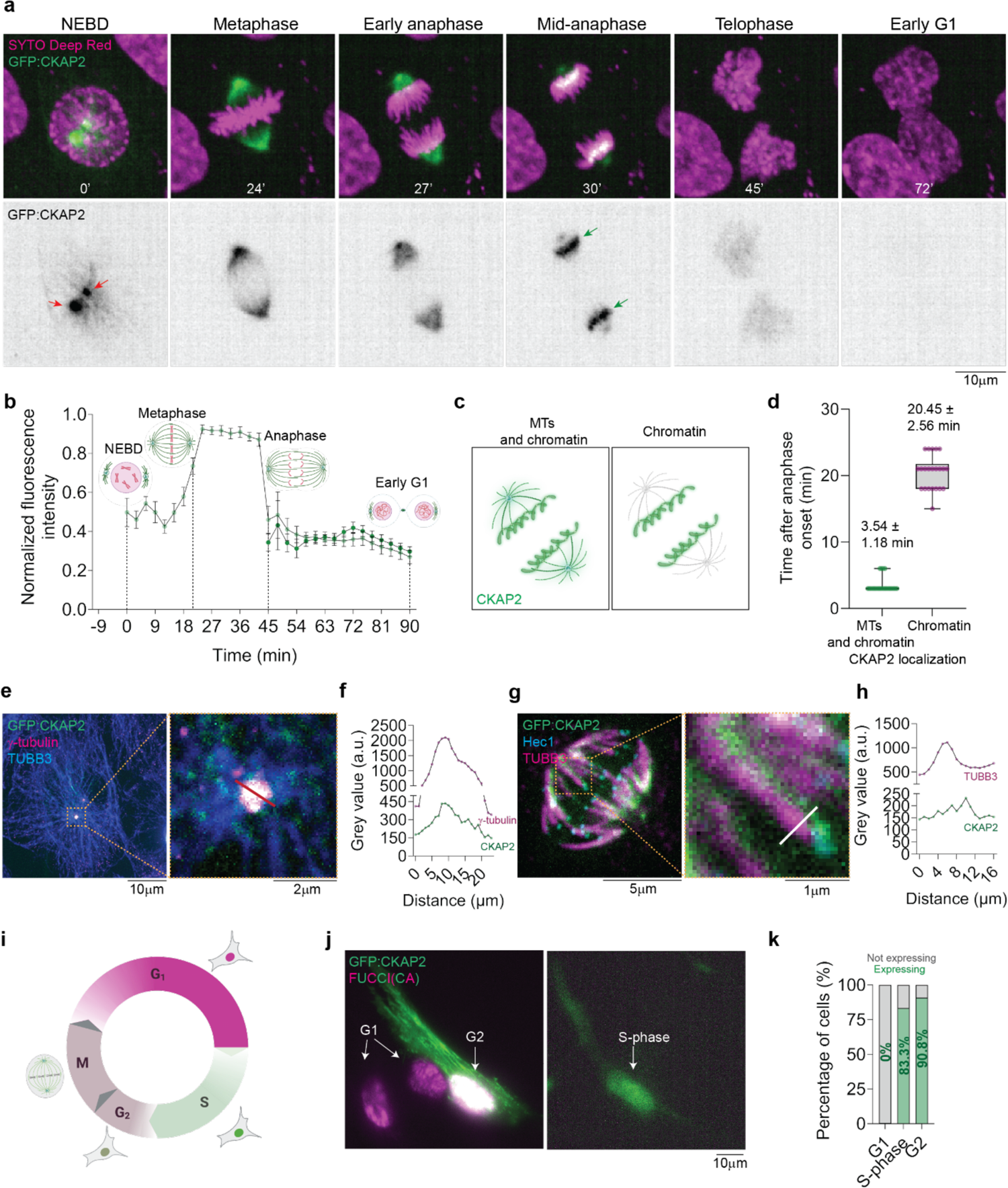
Cell cycle-dependent expression and subcellular localization of endogenously tagged CKAP2. **(a)** Representative time-lapse images of a GFP:CKAP2 knock-in RPE-1 cell undergoing a mitotic division. Note that CKAP2 initially localizes to the centrosomes at the onset of nuclear envelope breakdown (NEBD; red arrows). Note also that CKAP2 localization shifts from the spindle to chromatin after anaphase onset (green arrows) before being degraded. **(b)** Normalized fluorescence intensity of GFP:CKAP2 throughout mitosis (n = 22 cells) **(c)** Schematics of CKAP2 localization following anaphase. Upon anaphase onset, CKAP2 localization shifts to both microtubules and chromatin before being shifted entirely to the chromatin. **(d)** Quantification of time from anaphase onset until shift of CKAP2 localization to both microtubules (MTs) and chromatin and to chromatin only (n = 22 cells). **(e and f)** Representative image (e) and intensity profile plot (f) of a GFP:CKAP2 RPE-1 cell immunostained for γ-tubulin and β-tubulin (TUBB3). Note that CKAP2 co-localizes with γ-tubulin at the centrosome during interphase. **(g and h)** Representative image (g) and intensity profile plot (h) of a cold-treated GFP:CKAP2 RPE-1 cell immunostained for Hec1 and β-tubulin (TUBB3). Note that CKAP2 co-localizes with k-fibers during mitosis. **(i)** Schematics of FUCCI(CA) fluorescence patterns throughout the cell cycle. **(j)** Representative image of GFP:CKAP2 RPE-1 cells expressing FUCCI(CA) cell cycle marker. **(k)** Percentage of cells with observable CKAP2 expression at G1 (n = 207 cells), S-phase (n = 24 cells) and G2 (n = 219 cells). Time is shown in minutes. Measurements reported as average ± standard deviation.

Next, we set out to explore the dynamics of CKAP2 expression throughout interphase by transiently expressing the cell-cycle fluorescent marker FUCCI(CA) (27) in GFP:CKAP2 RPE-1 knock-in cells (Fig.1i-k). Briefly, FUCCI(CA) functions by monitoring the cell-cycle dependent proteolysis of Cdt1 and Geminin, and FUCCI(CA)-expressing cells display nuclear red fluorescence during G1, green fluorescence during S-phase and both red and green fluorescence during G2 and M-phase (27) (Fig. 1i). We detected that none of the cells found to be at G1 displayed visible CKAP2 fluorescence, whereas 83.3% of cells at S-phase and 90.8% of cells at G2 displayed clear CKAP2 fluorescence at cytoplasmic microtubules (Fig. 1j-k), indicating that CKAP2 expression is cell-cycle dependent. Overall, these first sets of experiments provide the first description of endogenously-labelled CKAP2 localization and dynamics throughout the cell cycle with high temporal resolution, and show the rapid patterns of CKAP2 redistribution from microtubules to chromatin upon mitotic exit.

### Cells lacking CKAP2 assemble morphologically-normal spindles but develop nuclear abnormalities and aneuploidy

Mis-regulation of CKAP2 is correlated with CIN and tumorigenesis, with increased CKAP2 levels being associated with tumor development and poor cancer prognosis (20, 23, 28). Similarly, RNAi-mediated knockdown of CKAP2 was found to interfere with spindle assembly and chromosome segregation in cells (17). We next sought to investigate the role of CKAP2 in cells by depleting CKAP2 expression using CRISPR-Cas9 knock-out (KO) (Supplementary Figure 2a). We isolated three KO clones from RPE-1 (C1, C2 and C3) and HT1080 cells (A2, B4 and B5), as confirmed by immunofluorescence, sequencing and Western Blot (Figure 2a; Supplementary Figure 2b-d), and we quantified mitotic and interphase phenotypes with immunofluorescence staining (Figure 2a-e; Supplementary Figure 3). Interestingly, we did not observe a significant impact of CKAP2 knock-out on mitotic phenotypes, as evidenced by no change in mitotic index (wild-type: between 3.9% to 4% mitotic cells; CKAP2-KO: between 1% to 5% mitotic cells) and in the proportion of multipolar spindles between wild-type and CKAP2-KO clones (wild-type: between 0% to 7% multipolar spindles; CKAP2-KO: 0% to 17% multipolar spindles; Figure 2b-c and Supplementary Figure 3b-c). On the other hand, severe nuclear abnormalities were frequently observed in all KO clones, including multinucleated/fragmented nuclei and “donut”-shaped nuclei, where a hollow surface that traverses the entire diameter of the nucleus is observed (Fig 2d-e; Supplementary Figure 3d-e), with only between 38% to 66% of CKAP2-KO clones displaying morphologically-normal nuclei as compared to between 95% to 98% in wild-type cells. Interestingly, in cells displaying donut-shaped nuclei, microtubules were often found to pass through the hole (Supplementary Figure 3f), suggesting that interphase microtubule architecture may be impacted in the absence of CKAP2.

**Figure 2.**
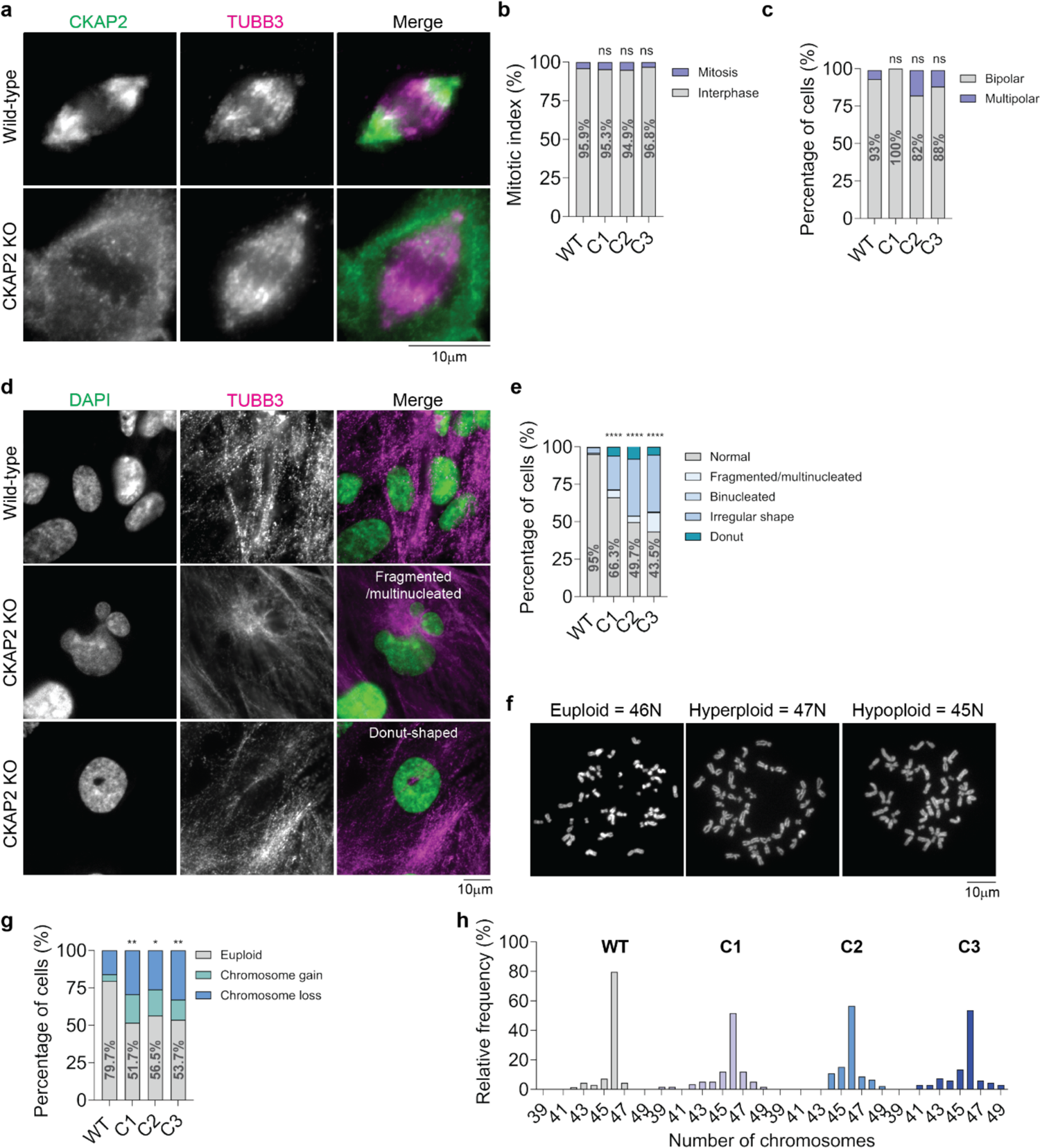
CKAP2 knock-out causes severe nuclear abnormalities and leads to aneuploidy. **(a and d)** Representative immunofluorescence images of wild-type (WT) and CKAP2 knock-out (KO) RPE-1 cells during mitosis **(a)** and interphase **(d)**. Note that KO cells display nuclear abnormalities such as multinucleated/fragmented nuclei and donut-shaped nuclei. **(b)** Quantification of percentage of WT (n = 1197 cells) and CKAP2 KO cells (C1 n = 680 cells; C2 n = 587 cells; C3 n = 630 cells) in interphase vs. mitosis (mitotic index). **(c and e)** Quantification of mitotic spindle **(c)** and interphase nuclei **(e)** abnormalities in WT (mitosis n = 15 cells; interphase n = 1152 cells) and CKAP2 KO clones C1 (mitosis n = 12 cells; interphase n = 650 cells), C2 (mitosis n = 17 cells; interphase n = 559 cells ****p < 0.0001) and C3 (mitosis n = 9 cells; interphase n = 607 cells ****p < 0.0001). **(f)** Representative images of metaphase spreads displaying euploidy, hyperploidy and hypoploidy. **(g)** Percentage of WT (n= 69 spreads) and CKAP2 KO cells (C1 n = 58 spreads **p = 0.0012; C2 n = 46 spreads *p = 0.0118; C3 n = 67 spreads **p = 0.0018) displaying chromosome gains and chromosome losses. **(h)** Frequency distribution of ploidy counts in WT (n= 69 spreads) and CKAP2 KO cells (C1 n = 58 spreads; C2 n = 46 spreads; C3 n = 67 spreads). Statistical significance relative to WT group. ns = non-significant at p β 0.05.

We next wondered whether the absence of CKAP2 impacts ploidy maintenance. Both RPE-1 and HT1080 cells are valuable models for ploidy studies, since wild-type cells are chromosomally stable, and display a modal chromosome number of 46 (ATCC.org), which we confirmed in our wild-type cells (Figure 2f-h; Supplementary Figure 4). Chromosome spreads revealed that two out of three HT1080 knock-out clones displayed near-tetraploid chromosome numbers (clones A2 and B5). In contrast, the third clone (B4) displayed a near-diploid chromosome set, albeit with more frequent chromosome gains and losses than wild-type cells (Supplementary Figure 4). All three RPE-1 knock-out clones displayed a near-diploid chromosome set and, similarly to HT1080 KO cells, harboured high levels of aneuploidy (Figure 2f-h). Altogether, our experiments demonstrate that CKAP2 is essential for genome stability, and the absence of CKAP2 causes elevated levels of aneuploidy and nuclear abnormalities. Further analysis was focused on the RPE-1 clones and the near diploid HT1080 clone to avoid any potential influence of tetraploidy on downstream analyses.

### Loss of CKAP2 leads to chromosome segregation errors and decreased microtubule growth rates

Several mitotic segregation errors have been shown to lead to aneuploidy in cells. Multipolar spindles that do not bipolarize prior to anaphase can lead to gross aneuploidy and cell cycle arrest/cell death (29); mitotic slippage and cytokinesis failure lead to tetraploidy (30) and anaphase lagging chromosomes can lead to single chromosome gains and losses and micronuclei formation (31, 32). These micronuclei can lead to abnormal chromosome rearrangements known as chromothripsis, which contribute to tumorigenesis (33). Thus, we next set out to explore how aneuploidy originates in CKAP2-KO cells by performing high-content screening live confocal imaging of wild-type and CKAP2-KO cells expressing CENPB:mEmerald (kinetochores), Tubulin:mRuby2 (microtubules) and stained with SYTO Deep Red (DNA) during cell division (Figure 3a; Supplementary Video 2). We found that CKAP2-KO cells displayed significantly higher proportions of chromosome segregation errors as compared to wild-type cells, with only between 52.8% to 73.9% of divisions being error-free in KO cells as opposed to between 81.8% to 94.8% in wild-type cells (Figure 3a-b; Supplementary Figure 5a-b). Consistent with our immunofluorescence experiments (Figure 2; Supplementary Figure 3), the majority of cells assembled morphologically normal bipolar spindles similarly to wild-type cells (Figure 3b; Supplementary Figure 5b), with a small proportion of CKAP2-KO cells displaying multipolar spindles (Figure 3a bottom right inset; between 2.2% to 2.5%). However, CKAP2-KO cells showed severe chromosome segregation errors, including lagging chromosomes that often resulted in micronucleus formation, chromatin bridges and chromosome misalignments (Figure 3a-c; Supplementary Figure 5a-d). Importantly, loss of CKAP2 had no impact on mitosis duration and bipolar spindle length in RPE-1 cells and caused a negligible increase of mitosis duration and decrease in spindle length in HT1080 cells (Figure 3d-e; Supplementary Figure 5e-g), suggesting that the segregation errors originated in the absence of CKAP2 may be a result of more subtle alterations in microtubule dynamics rather than gross defects in spindle assembly/chromosome congression and multipolarity.

**Figure 3.**
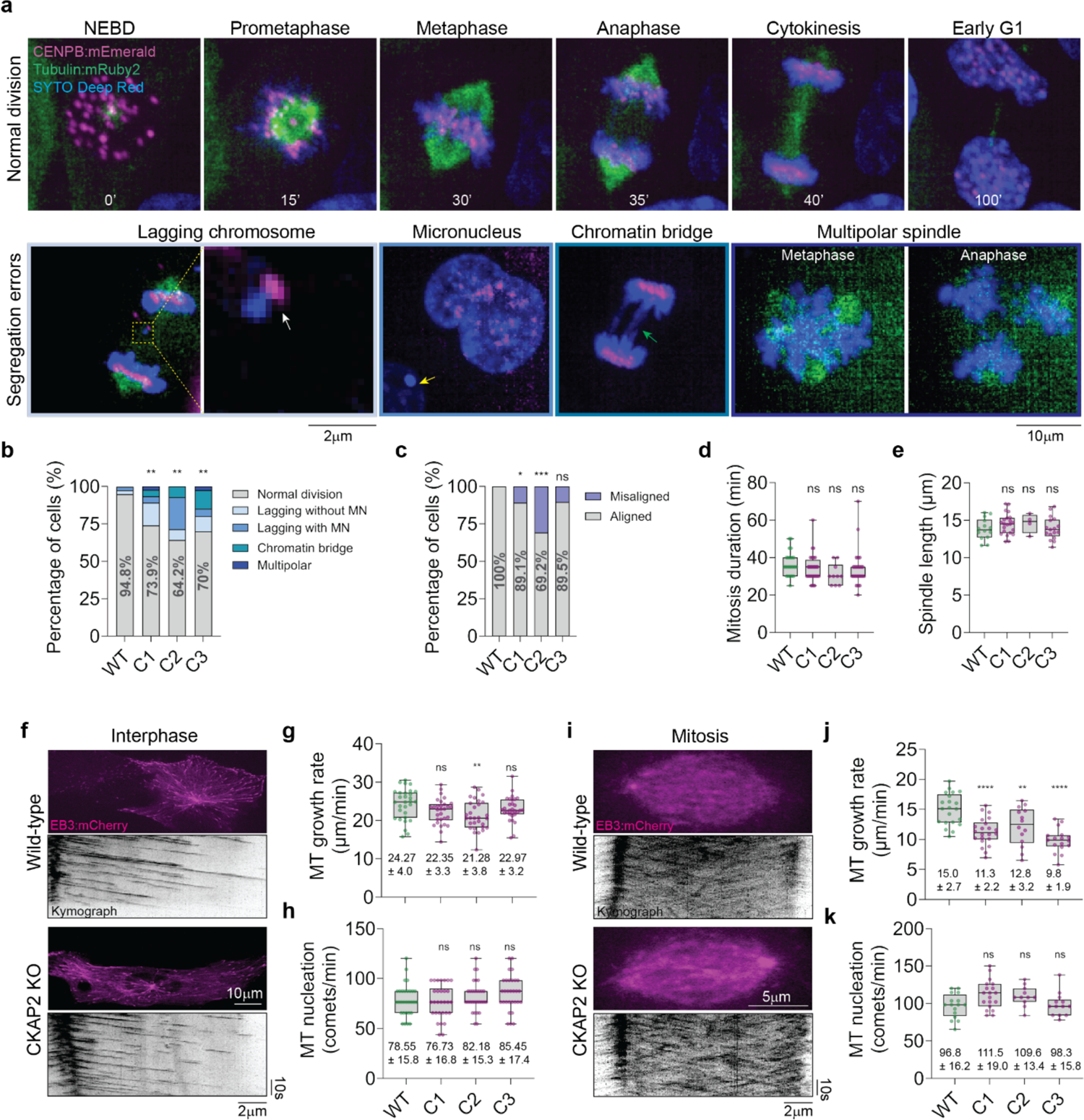
CKAP2 knock-out impacts chromosome segregation and microtubule dynamics. **(a)** Representative time-lapse images of a wild-type (WT) RPE-1 cell co-expressing CENPB:mEmerald (magenta), Tubulin:mRuby2 (green) and stained with SYTO Deep Red DNA label (blue) undergoing a mitotic division (top), and examples of chromosome segregation errors observed in CKAP2 KO cells (bottom). CKAP2 KO cells frequently displayed anaphase lagging chromosomes (white arrow), micronuclei formation (yellow arrow) and chromatin bridges (green arrow), and occasionally displayed multipolar spindles. **(b)** Quantification of chromosome segregation errors in WT (n = 39 cells) and KO cells (C1 n = 46 cells **p = 0.0094; C2 n = 14 cells **p = 0.0037; C3 n = 40 cells **p = 0.0038). **(c)** Percentage of cells displaying aligned vs. misaligned chromosomes at anaphase onset in WT (n = 34 cells) and KO cells (C1 n = 46 cells *p = 0.0471; C2 n = 13 cells ***p = 0.0007; C3 n = 38 cells). **(d)** Quantification of mitosis duration measured as the time from nuclear envelope breakdown (NEBD) to the first frame of anaphase onset in WT (36.74 ± 6.5 min; n = 23 cells) and KO cells (C1 34.22 ± 7.2 min n = 32 cells; C2 31 ± 5.7 min n = 10 cells; C3 33.9 ± 9.3 min n = 31 cells). **(e)** Quantification of metaphase spindle length measured at the last frame prior to anaphase onset in WT (13. 82 ± 1.4 µm; n = 14 cells) and KO cells (C1 14.47 ± 1.4 µm n = 22 cells; C2 14.61 ± 1.3 µm n = 4 cells; C3 13.91 ± 1.5 µm n = 17 cells). **(f)** Representative snapshots (top) and kymographs (bottom) of wild-type (WT) and CKAP2 knock-out (KO) interphase cells expressing EB3:mCherry (magenta). **(g and h)** Quantification of microtubule growth rates **(g)** and nucleation from centrosomes **(h)** in WT (n = 30 cells) and KO interphase cells (C1 n = 30 cells; C2 n = 30 cells; C3 n = 30 cells). **(i)** Representative snapshots (top) and kymographs (bottom) of wild-type (WT) and CKAP2 knock-out (KO) mitotic cells expressing EB3:mCherry (magenta). **(j and k)** Quantification of microtubule growth rates **(g)** and nucleation from centrosomes **(h)** in WT (nucleation: 96.82 ± 16.2 comets/min n = 16 cells; growth: 15 ± 2.6 µm/min n = 21 cells) and KO mitotic cells (C1 nucleation: n = 19 cells; growth: n = 22 cells ****p < 0.0001; C2 nucleation: n = 11 cells; growth: n = 14 cells **p = 0.0089; C3 nucleation: n = 13 cells; growth: n = 17 cells ****p < 0.0001). Time is shown in minutes. Statistical significance relative to WT group. ns = non-significant at p ≥ 0.05. Measurements reported as average ± standard deviation.

Microtubules undergo continuous cycles of growth and shrinkage, a property known as dynamic instability that is key for cellular processes ranging from cell division, organelle transport and neuronal communication (7, 34). Increased microtubule growth rates have been observed in chromosomally unstable cell lines and are thought to contribute to the emergence of aneuploidy (5). Our recent *in vitro* reconstitution experiments have demonstrated that CKAP2 increases microtubule nucleation and growth (24). Therefore, we next explored whether microtubule nucleation and growth dynamics are affected in cells lacking CKAP2.

For this purpose, we performed live imaging of cells expressing fluorescently labelled plus-end directed protein EB3 (EB3:mCherry (35) or EB3:tdStayGold (36)) and measured microtubule nucleation rates (as quantified by the number of newly assembled comets from centrosomes per minute) and microtubule growth rates with kymograph plots (see Methods for more details; Figure 3f-k; Supplementary Figure 5h-m). We found no difference in nucleation rates between wild-type and CKAP2-KO cells both during interphase and mitosis (Figure 3f-k; Supplementary Figure 5h-m). However, we detected a significant decline of about 20% (from 15 um/min to 11.1 um/min) in microtubule growth rates in CKAP2-KO cells compared to wild-type cells during mitosis, but not during interphase in RPE-1 cells (Figure 3f-k; Supplementary Figure 5h-m). Interestingly, by performing a cold shock approach that allows for the visualization of stable kinetochore-microtubule attachments (see Methods for details), we found that CKAP2-KO cells displayed a significantly increased percentage of merotelic attachments – wherein a single kinetochore is attached to microtubules emanating from both spindle poles - and unattached kinetochores per cell, as compared to wild-type cells (wild-type: 26.67% vs. KO: 93.33% of KO cells displaying at least one mis-attachment), which may contribute to chromosome mis-segregation (Supplementary Figure 6a-c). These results correlate with the high levels of segregation errors observed in CKAP2 KO cells (Figure 3) and are consistent with the notion that even a single merotelically-attached kinetochore is sufficient to cause segregation errors (31). Our results demonstrate that the absence of CKAP2 negatively impacts chromosome segregation fidelity, disrupts microtubule growth dynamics during mitosis and causes the kinetochore-microtubule mis-attachments.

### CKAP2 expression alters chromosome segregation and microtubule growth dynamics

Previous reports have shown that CKAP2 is upregulated in various human carcinomas (20, 23, 37–39), and overexpression of CKAP2 in cells impairs nuclear integrity and ploidy maintenance (18). Thus, we next explored whether exogenous CKAP2 expression in wild-type and CKAP2-KO cells could alter chromosome segregation dynamics. For that purpose, we took advantage of a doxycycline-inducible expression system to control the levels of CKAP2 expression in cells (see Methods for more details). Live cell imaging experiments demonstrated that wild-type cells expressing CKAP2:mGL displayed an increased proportion of chromosome segregation errors, with only 69.7% of those cells undergoing error-free divisions, as opposed to 86.56% in wild-type cells not expressing CKAP2:mGL (Figure 4a-c). The levels of chromosome mis-segregation observed in wild-type cells expressing CKAP2:mGL were similar to those found in CKAP2-KO cells (64.78% of error-free divisions in CKAP2-KO cells; Figure 4a-c), indicating that either excessive expression or an absence of CKAP2 can impair chromosome segregation to a similar extent. Interestingly, when CKAP2:mGL was exogenously expressed in CKAP2-KO cells, the proportion of chromosome segregation errors was identical to that of wild-type cells, with 84.2% of KO cells expressing CKAP2:mGL displaying error-free divisions (Figure 4a-c). These experiments indicate that chromosome segregation fidelity can be rescued in CKAP2-KO cells by exogenous expression of CKAP2.

**Figure 4.**
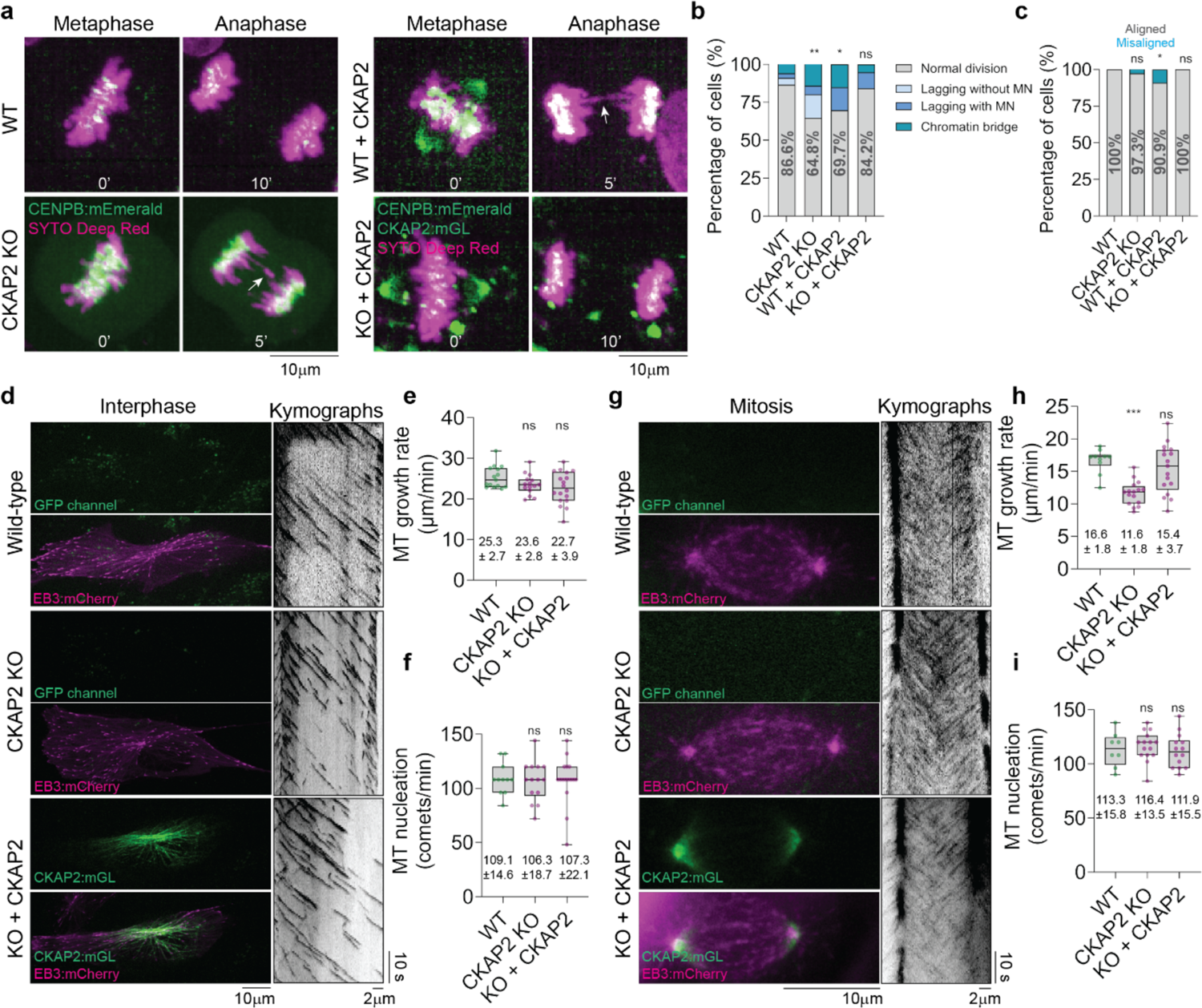
CKAP2 expression alters chromosome segregation and spindle microtubule growth dynamics. **(a)** Representative time-lapse images of wild-type (WT), CKAP2 knock-out (KO), WT expressing CKAP2:mGL and CKAP2 KO expressing CKAP2:mGL RPE-1 cell undergoing a mitotic division. **(b)** Quantification of chromosome segregation errors in WT (n = 67 cells), KO cells (n = 71 cells; **p = 0.0030), WT cells expressing CKAP2:mGL (n = 33 cells, *p = 0.0432) and KO cells expressing CKAP2:mGL (n = 19 cells). **(c)** Percentage of cells displaying aligned vs. misaligned chromosomes at anaphase onset in WT (n = 67 cells), KO cells (n = 71 cells), WT cells expressing CKAP2:mGL (n = 33 cells, *p = 0.0122) and KO cells expressing CKAP2:mGL (n = 17 cells). **(d)** Representative snapshots (left) and kymographs (right) of wild-type (WT), CKAP2 KO cells and CKAP2 KO cells expressing CKAP2:mGL and EB3:mCherry (magenta) during interphase. **(e and f)** Quantification of growth rates **(e)** and nucleation from centrosomes **(f)** in WT (nucleation: n = 11 cells; growth: n = 15 cells), KO cells (nucleation: n = 14 cells; growth: n = 15 cells) and KO cells expressing CKAP2:mGL (nucleation: n = 16 cells; growth: n = 18 cells) during interphase. **(g)** Representative snapshots (left) and kymographs (right) of wild-type (WT), CKAP2 KO cells and CKAP2 KO cells expressing CKAP2:mGL and EB3:mCherry (magenta) during mitosis. **(h and i)** Quantification of microtubule growth rates **(h)** and nucleation from centrosomes **(i)** in WT (nucleation: n = 8 cells; growth: n = 10 cells), KO cells (nucleation: n = 15 cells; growth: n = 16 cells ***p = 0.0002) and KO cells expressing CKAP2:mGL (nucleation: n = 14 cells; growth: n = 17 cells) during mitosis. Time is shown in minutes. Statistical significance relative to WT group. ns = non-significant at p ≥ 0.05. Measurements reported as average ± standard deviation.

Moreover, similarly to our previous results (Figure 3f-k), plus-end microtubule tracking experiments revealed that CKAP2-KO cells display reduced microtubule growth rates (11.66 ± 1.83 µm/min) as compared to wild-type cells during mitosis (16.66 ± 1.87 µm/min) with no observable changes on microtubule nucleation and interphase growth rates (Figure 4d-ij). When CKAP2:mGL was expressed in KO cells, microtubule growth rates during mitosis normalized to that of wild-type cells (KO+CKAP2: 15.40 ± 3.73 µm/min), whereas microtubule nucleation and interphase growth rates remained unchanged (Figure 4d-i), indicating that CKAP2 expression is sufficient to rescue disrupted microtubule growth dynamics during mitosis caused by CKAP2 knock-out. Altogether, our results demonstrate that CKAP2 acts to accelerate microtubule growth during mitosis in cells, consistent with our measurements *in vitro* (24), and contributes to proper chromosome segregation and ploidy maintenance likely by ensuring the establishment of proper kinetochore-microtubule attachments.

## Discussion

Our CRISPR-Cas9 knock-in live cell imaging experiments revealed the dynamics of endogenously-labelled CKAP2 expression and localization throughout the cell cycle with high temporal resolution (Figure 1). We demonstrate that CKAP2 expression initiates during the S- to G2-phase transition, reaches its peak during metaphase and further declines at early G1 of the following cell cycle, consistent with the notion that CKAP2 degradation is dependent on the anaphase promoting complex/cyclosome (APC/C) targeting CKAP2 for degradation upon mitotic exit (14, 15). Our live cell experiments reveal a shift in CKAP2 from microtubules to chromatin after the onset of anaphase, which is consistent with previous reports (14, 15, 25) and is thought to be dependent on CKAP2 phosphorylation at Ser627 (26). Overall, our experiments enabled us to assess the precise timescale of CKAP2 dynamics further, revealing that the microtubule-to-chromatin shift occurs sequentially – first localizing to both microtubules and chromatin before being shifted entirely to the chromatin - and rapidly within ∼20 min of anaphase onset. Why CKAP2 localization changes from microtubules to chromatin is unknown but may indicate a role of CKAP2 on the regulation of nuclear envelope reformation and/or chromatin integrity. Consistent with this idea, our CRISPR-Cas9 knock-out experiments demonstrate that CKAP2 loss causes severe nuclear abnormalities (Figure 2 and Supplementary Figure 3).

How CKAP2 loss impairs chromatin integrity is unclear. However, we find that CKAP2-KO cells frequently display abnormal anaphase movement, in which chromosomes begin to separate but quickly reverse their direction, forming a U-shaped chromosome mass that appears to contribute to donut-shaped nuclear formation (Supplementary Figure 7). Moreover, some CKAP2-KO clones display reduced speed of chromosome separation (Supplementary Figure 7). Similar anaphase movement defects have been reported to occur in HeLa-Kyoto cells expressing mutants of Kif22, a mitotic kinesin motor (40). These abnormal anaphase movements are thought to result from failure of Kif22 inactivation at anaphase onset, thereby disrupting force balance at anaphase (40). Donut-shaped nuclei have also been observed in cells overexpressing TPX2, and these cells were found to house a single centrosome inside their hollow surface (12), similar to our observations in CKAP2-KO cells (Supplementary Figure 3). It is possible that altered microtubule dynamics could impact midzone microtubule growth, which is crucial for anaphase chromosome separation (41), and this could thus account for the anaphase movement defects and abnormal nuclei formation, as observed in cells overexpressing TPX2 (11) and cells lacking CKAP2 in our study. Alternatively, the shift in CKAP2 localization from microtubules to chromatin may suggest that CKAP2 has additional microtubule-independent roles on chromatin structure and integrity, as suggested by a previous report showing that CKAP2 knockdown impairs nuclear lamina distribution (17), however, the mechanistic basis of how CKAP2 maintains nuclear integrity remains to be further explored.

Our CRISPR-Cas9 knock-out experiments also revealed that loss of CKAP2 leads to aneuploidy, with 4 out of 6 CKAP2-KO clones displaying near-diploid chromosome numbers, although with high levels of aneuploidy, and the remaining 2 clones containing a near-tetraploid chromosome set (Figure 2 and Supplementary Figure 3). Importantly, our live imaging experiments revealed that genome doubling events (cytokinesis failure/mitotic slippage) were very rare in CKAP2-KO cells (Figure 3 and Supplementary Figure 5), suggesting that the near-tetraploid phenotype detected in the HT1080 clones A2 and B5 may have originated from early tetraploidization events taking place before or soon after knock-out, rather than as a direct result of CKAP2 loss.

We find that CKAP2 directly regulates microtubule dynamics, and cells lacking CKAP2 develop high levels of chromosome segregation errors due to a reduction in microtubule growth rates during mitosis. The slowdown in microtubule growth rates by about 20% likely causes the accumulation of mis-attached kinetochores that result in chromosome mis-segregation (Figures 3 and 4). Previous work has shown that knockdown of the MAP chTOG/XMAP215/CKAP5 reduces microtubule growth rates in cells (5, 42, 43), promoting an average reduction of 1.8 μm/min in chromosomally stable cell lines (5). Our results demonstrate a substantially greater influence of CKAP2 KO on reducing microtubule growth rates as compared to chTOG/XMAP215/CKAP5, with an overall reduction of between 2.82 to 5.16 μm/min in mitotic CKAP2-KO cells, thus indicating that an excessive decline in microtubule growth rates is a contributing factor for the development of chromosome mis-segregation. Moreover, altered kinetochore-microtubule dynamics driven by the absence of CKAP2 in our experiments is not a result of gross spindle architecture defects, as evidenced by the low proportions of multipolar spindles in CKAP2 KO cells (Figures 2 and 3; Supplementary Figures 3 and 5), and is thus most likely a direct impact of reduced microtubule growth rates, unlike the impact of increased microtubule growth rates (5).

Here we also demonstrate that microtubule nucleation is unaffected upon loss of CKAP2 in cells. This is in stark contrast to our previous findings that CKAP2 dramatically increases microtubule nucleation *in vitro* (24), and suggest that CKAP2 may not have a major role on centrosomal microtubule nucleation in cells, or that other centrosomal components could compensate for the lack of CKAP2 *in vivo*, given the plethora of MAPs that induce microtubule nucleation (44) present at the centrosome (45). Consistent with this notion, recent work has shown that the MAPs chTOG, CLASP1, CAMSAP and TPX2 can all independently nucleate microtubules in the absence of γ-tubulin in HCT116 cells (46), and it is thus possible that any those factors may overcome the absence of CKAP2 and recover nucleation rates in KO cells. This is also in line with our findings that CKAP2 loss has no substantial impact upon spindle size (Figure 3 and Supplementary Figure 5), as opposed to previous reports showing a reduction in spindle length upon depletion of the microtubule polymerase XMAP215 (47–49). These differences may be attributed to CKAP2 acting predominantly as a microtubule growth factor in cells, in contrast to XMAP215, which has both microtubule growth (13) and nucleation (50) functions and is also consistent with the notion that microtubule nucleation is essential for spindle length determination (51). Nevertheless, our data shows that the regulatory role of CKAP2 on microtubule growth is independent of its role as a microtubule nucleator, as evidenced by the decline in microtubule growth rates in KO cells regardless of unchanged nucleation rates.

Chromosome mis-segregation and CIN favours the proliferation of cells with tumorigenic potential and is considered a hallmark of cancer (52). Previous work has shown that CKAP2 is involved in the maintenance of genome integrity (16–19), and our results further build upon the current knowledge of CKAP2’s functions by providing mechanistic evidence that CKAP2 ensures accurate chromosome segregation by regulating microtubule dynamics, specifically microtubule growth at the spindle. Overall, our results reveal the essential role of CKAP2 on microtubule dynamics and provide a mechanistic explanation for the oncogenic potential of CKAP2 mis-regulation.

## Methods

### Cell Culture

HT-1080 (CCL-121 – ATCC) cells were obtained from Synthego, and hTERT RPE-1 cells were a gift from Arnold Hayer (McGill University). Cells were cultured in either DMEM (HT1080) or DMEM/F-12 (RPE-1) with high glucose and sodium pyruvate supplemented with 10% fetal bovine serum, 1% Penicillin/Streptomycin and 10 μg of Hygromycin (DMEM/F-12 only). Cells were cultured at 5% CO_2_ and 37°C.

### Generation of CKAP2 knock-in and knock-out cells by CRISPR/Cas9

For CRISPR/Cas9 knock-in, Cas9 Nuclease 2-NLS, sgRNAs targeting exon 4 of CKAP2 and a template DNA generated by PCR containing a GFP and Blasticidin resistance sequence were used. For CRISPR/Cas9 knock-out, Cas9 Nuclease 2-NLS and a sgRNAs targeting exon 4 of CKAP2 were used. For both knock-in and knock-out, RNP complexes were formed in Nucleofector solution at a 6:1 ratio (sgRNA:Cas9) and immediately transfected by Nucleofection into HT-1080 cells using the Amaxa SF Cell Line 4D-Nucleofector X kit S (Lonza, PBC2-00675) and either program FF-113 (HT1080) or EA-104 (RPE-1) on the 4D-Nucleofector X unit. Cells were harvested, and 2 x 10^5^ cells were added to the RNP complex mix in a 16-well Nucleocuvette. After nucleofection, the transfected mixes were resuspended in warm complete DMEM or DMEM/F-12, plated in 24-well plates, and media was changed 24h after transfection.

### Stable cell line generation

Stable HT1080 knock-out cell lines were isolated from the edited pools by limiting dilution in 96-well plates for 5 weeks. Briefly, the edited pool was diluted to a working concentration of 1.8 cells per well in warm complete DMEM. This dilution was seeded on 96-well plates and incubated under regular conditions. The plates were monitored weekly for colony formation until 80% well confluency was reached (5-weeks), then single cell colonies were expanded in 24-well plates. Stable RPE-1 knock-out clones were isolated from edited pools using a DispenCell® Single-Cell Dispenser according to manufacturer’s instructions. After expansion, clones were sequenced and tested by western blot and immunofluorescence for the absence of CKAP2 protein staining. For stable knock-in cell line isolation, cells were treated with Blasticidin (HT1080 - 0.5μg/mL; RPE-1 – 1.6µg/mL) and confirmation of GFP:CKAP2 expression was done with confocal microscopy. For the FUCCI(CA) experiment in Figure 1i-k, a single RPE-1 GFP:CKAP2 knock-in clone was isolated from the knock-in pool using DispenCell® Single-Cell Dispenser and was used for the experiment.

### Sequencing

To validate the genomic CRISPR edit presence, stable KO and KI cell lines cells were grown on 24-well plates for 48 h and their DNA was collected with QuickExtract Solution (Mandel Scientific LGN-QE0905T). PCR was conducted with primers targeting a region on exon 4 for human CKAP2. PCR mixes containing 1 U of PfuX7 polymerase 40, 1x reaction buffer, 10 mM dNTPs, 1 µL of QuickExtract DNA, 10 mM primers and nuclease free H2O (up to 50 µL) were ran on a DNAEngine TETRAD2 Peltier Thermal cycler (MJ Research) with the following program: initial denaturation at 98°C for 10 min followed by 40 cycles of denaturation at 98°C for 30 s, annealing at 58°C for 30 s and elongation at 72°C for 1 min, final elongation at 72°C for 7 min and a 4°C hold. PCR products were visualized on a 1 % agarose gel and sent for Sanger Sequencing (Genome Quebec) with a nested sequencing primer. Sequencing data was analyzed with Synthego’s ICE software.

### Metaphase spreads

Cells were seeded and grown in complete culture media for 24h, then the media was washed off and replaced with complete media containing 100 µM Monastrol for 4h. Cells were then washed twice with PBS and trypsinized to collect a cell suspension. Subsequently, the cells were pelleted and resuspended in a hypotonic solution (0.056M KCl) and incubated at 37°C for 30 min before being pelleted and fixed in a methanol: glacial acetic acid (3:1) solution. Fixed cells were dropped into glass slides and stained with DAPI. Imaging was performed on a Zeiss Axio Observer 7 fitted with a Colibri and Alpha Plan-Apo 100x/1.46Oil objective.

### Cold shock

For assessment of kinetochore-microtubule attachments in Supplementary Figure 6, cells were exposed to a brief cold-shock treatment that serves to depolymerize less stable non-kinetochore microtubules, allowing for the clear visualization of kinetochore-microtubule attachments. Briefly, cells were treated with 50 ng/mL Nocodazole for 4h to induce a metaphase arrest, followed by a wash out of Nocodazole in pre-warmed DMEM and incubation at 37°C and 5% CO_2_ for 13 min for spindle re-assembly. Cells were then exposed to cold treatment on ice for 12 min, followed by fixation for immunofluorescence staining.

### Immunofluorescence

Cells were either cultured on glass coverslips or on black 96-well glass-bottom PerkinElmer plates until 70% confluency, and then washed twice with PBS and fixed with −20 °C methanol. Subsequently, cells were incubated with fresh blocking solution (either 5% BSA, 0.05% Tween-20 in 1x PBS or 3% BSA in 1x PBS) overnight at 4°C, followed by incubation with primary antibodies diluted in blocking solution overnight at 4°C. Next, cells were washed 2 times with blocking solution and incubated with secondary antibodies for 2h at room temperature. Then, cells were washed 3 times with PBS and incubated with DAPI (1µg/mL) for 15 min at room temperature, finally, the coverslips were washed 2 times with PBS and mounted on slides with FluorSave. Once the FluorSave had polymerized and cured for 24h, cells were visualized by spinning disk microscopy on a Quorum Diskovery platform installed on a Leica DMi8 inverted microscope. This system consists of an HCX PL APO 63x/1.4 NA oil objective, a DISKOVERY multi-modal imaging system (Spectral) with a multi-point confocal 50 µm pinhole spinning disk and dual iXon Ultra 512×512 EMCCD (Andor) cameras for simultaneous imaging, ASI three axis motorized stage controller, and MCL Nano-view piezo stage, 488 nm, 561 nm and 647 nm solid state OPSL lasers linked to a Borealis beam conditioning unit. Image acquisition and microscope control was executed using MetaMorph (Molecular Devices). For experiment in Supplementary Figure 1c-d, imaging was performed on a Zeiss Axio Observer 7 fitted with a TIRF and Alpha Plan-Apo 100x/1.46Oil objective. For experiment in Supplementary Figure 2b, cells were imaged on a spinning disk confocal Opera Phenix Plus High-Content Screening System containing a 63x/1.15 NA water objective. Primary antibodies used were rabbit polyclonal anti-CKAP2 (Proteintech, 25486-1-AP, 1:200), rabbit polyclonal anti-TUBB3 (BioLegend, 802001, 1:200), mouse monoclonal anti-Hec1 (BioLegend, sc-515550, 1:25), mouse monoclonal anti-γ−tubulin (Sigma Aldrich, T5326 1:1000), mouse monoclonal anti-β-tubulin TUB 2.1 (Sigma Aldrich, T4026, 1:200). AlexaFluor secondary antibodies (488nm and 548nm) were purchased from Invitrogen and used at 1:1000 concentrations.

### Western Blotting

Cells were cultured in 6-well plates until desired confluency, quickly washed in PBS, and directly lysed in RIPA buffer (150 mM NaCl, 1.0% Triton X-100, 0.5% sodium deoxycholate, 0.1% SDS, 50 mM Tris, pH 8.0). Proteins were gently homogenized and mixed 1:2 with Laemmli loading buffer, denatured for 5 min at 95°C and resolved by SDS-Page electrophoresis, transferred to ethanol activated PVDF membranes and blocked in 5% milk and 0.1% Tween-20 for 30 min. Primary incubation was done overnight at 4 °C with rabbit anti-CKAP2 1:2000 or mouse anti-GAPDH (BioLegend, 649201, 1:3000. For secondary antibodies, HRP-linked anti-rabbit or anti-mouse were used, and the HRP signal was visualized with ECL Western Blotting Detection reagents (ABM) as per the manufacturer’s instructions. Imaging was performed with a Chemidoc MP Imaging System (BioRad).

### Plasmid transfection

Plasmid DNA was transformed into NEB Turbo competent *E. coli* and midipreps were prepared using ZymoPURE II Plasmid Midiprep kit (ZYMO Research, D4201) according to manufacturer’s instructions. Plasmids used were H2B:mCerulean (Addgene, 55374), Tubulin:mRuby2 (Addgene, 55915), EB3:tdStayGold (36) (gift from Atsushi Miyawaki, RIKEN Japan), EB3:mCherry (35) (gift from Alyson Fournier, McGill University), CENPB:mEmerald (Addgene, 54037) and FUCCI(CA) (Addgene, 153521). For EB3:tdStayGold, EB3:mCherry and FUCCI(CA), transfection was performed using Xfect Transfection Reagent (Takara Bio, 631317) with 5 μg of DNA following manufacturer’s instructions and cells were imaged 24h post-transfection. For H2B:mCerulean and Tubulin:mRuby2 co-transfection, as well as CENPB:mEmerald and Tubulin:mRuby2 co-transfection, cells were transfected using Nucleofection with an Amaxa SF Cell Line 4D-Nucleofector X kit S (Lonza) and program FF-113 (HT1080) or EA-104 (RPE-1) on the 4D-Nucleofector X unit with a total of 1.3 μg of DNA and imaged 48h post-transfection.

### Doxycycline-inducible CKAP2:mGL expression

For experiments involving exogenous CKAP2:mGL expression, the Tet-On® 3G Inducible Expression System kit was used (Takara Bio; 631167). RPE-1 wild-type and CKAP2-KO cells were transfected with a pEF1α-TET3G plasmid for expression of a Tet-On 3G transactivator protein, and stable cell lines were then generated by antibiotic selection using 500µg/mL of G418. pEF1α-TET3G-stable cell lines were then transfected with a pTRE3G-CKAP2:mGL plasmid containing the P_TRE3G_ promoter activatable upon doxycycline exposure and cultured in regular DMEM/F-12 media supplemented with Tet-Approved FBS free of tetracycline contaminants (Takara Bio; 631105). Prior to experiments, transfected cells were exposed to 8ng/mL of doxycycline (Thermo Scientific; J60422-06) for ≥12h for induction of CKAP2:mGL expression.

### Live cell imaging

For GFP:CKAP2 knock-in live imaging in Fig. 1 and chromosome segregation dynamics live imaging in Fig 3 and 4, cells were cultured on black 24- or 96-well glass-bottom PerkinElmer plates in complete culture media at 37 °C and 5% CO_2_ and grown until 70% confluency. For experiments in Figures 1, 3, 4 and Supplementary Figure 1b cells were treated with 500 nM SYTO Deep Red for live DNA staining 20 min prior to imaging. Cells were imaged on spinning disk confocal Opera Phenix Plus High-Content Screening System containing a 63x/1.15 NA water objective at 37 °C and 5% CO_2_ for 4h at either 10 min (Supplementary Figure 1b), 3 min (Figure 1) or 5 min (Figure 3, 4 and Supplementary Figure 5a-g) interval acquisitions. For EB3:tdStayGold and EB3:mCherry live imaging in Figure 4 and Supplementary Figure 5h-m, cells were cultured on glass bottom FluoroDishes (WPI) in complete culture media at 37°C and 5% CO_2_ and grown until 70% confluency. Cells were treated with 500 nM SYTO Deep Red for live DNA staining (ThermoFischer) 20 min prior to imaging and for mitosis analysis, cells were treated with 50 ng/mL Nocodazole for 4h for metaphase arrest and then released into fresh media for 30 min prior to imaging. Images were acquired on a Quorum WaveFX-X1 spinning disk confocal system, on a Leica DMI6000B inverted microscope at 37°C and 5% CO_2_ equipped with a 63x/1.4-0.6 NA oil lens, at 500 ms for 1 min with an exposure time of 500 ms and microscope control was executed using MetaMorph (Molecular Devices).

### Image and data analysis

All images were processed and analyzed using Fiji (ImageJ). Microtubule dynamics in interphase and mitotic cells were analyzed using kymographs. Microtubule growth rates were measured by manually drawing lines on kymographs and measuring the slope of growth. Microtubule nucleation rates were measured by counting the number of EB3 comets emerging from centrosomes over the course of 2.5 seconds and then adjusting to the duration of the movie (60 s) and are reported in units of new comets emerging from the centrosome per minute. Speed of chromosome separation was quantified as (D_2_ – D_1_)/5, where D is the distance between the two chromosome masses at the first (D1) and second (D2) frames of anaphase (time interval of 5 minutes), and reported as μm per minute. All statistical analyses and data plotting were performed using GraphPad Prism 9 (www.graphpad.com). In the box plots, the center line represents the median, the bounds of the box represent upper and lower quartiles, the whiskers represent minimum and maximum values, and dots represent independent measurements. For numerical data, Shapiro–Wilk normality tests were applied and either unpaired two-tailed t tests or unpaired two-tailed Mann–Whitney U tests were applied. For categorical data fitting contingency tables, Fischer’s exact test was applied. Figures were assembled using Adobe Illustrator and illustrations in Supplementary Figures 1a and 2a were assembled using BioRender.

## Acknowledgements

This work was funded by grants from the Canadian Institutes of Health Research (PJT-156193) and Natural Sciences and Engineering Research Council of Canada (NSERC; RGPIN-2017-04649). LMGP is supported by an NSERC Postdoctoral Fellowship, AALJ was supported by a Mitacs Globalink Graduate Fellowship and TSM was supported by Doctoral Scholarships from Fonds de Recherche du Québec Santé and the Centre de Recherche en Biologie Structurale. We thank the members of the Bechstedt and Brouhard’s laboratories for constructive discussions on the project. We also thank the McGill Advanced Bioimaging Facility and the McGill Imaging and Molecular Biology Platform for training and technical support with microscopy experiments and the CRISPR/IPSC platform at the Montréal Neurological Institute for their help with the generation of CRISPR cell lines.

## Author contributions

LMGP: Data curation, formal analysis, investigation, methodology, validation, visualization, writing – original draft, writing – review and editing.

AALJ: Data curation, formal analysis, investigation, methodology, validation, visualization. TSM: Formal analysis, visualization.

SB: Conceptualization, funding acquisition, investigation, methodology, project administration, supervision, writing - original draft, writing – review and editing

## Supplementary Figures

**Supplementary Figure 1.**
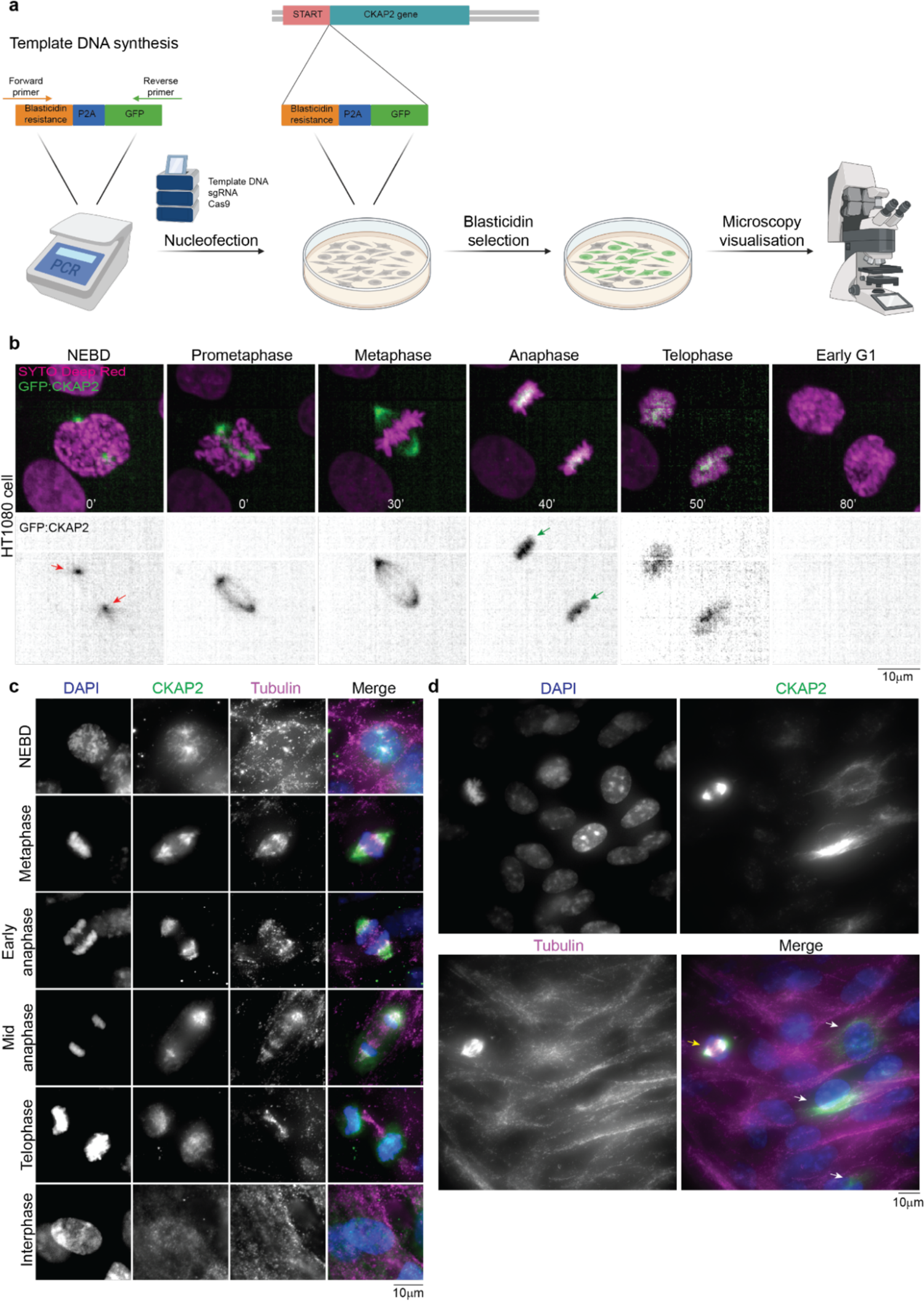
Subcellular localization of CKAP2. **(a)** Illustration detailing the experimental design for CRISPR-Cas9 knock-in. An N-terminally tagged GFP sequence linked to a Blasticidin resistance gene sequence were amplified by PCR and transfected into RPE-1 and HT1080 cells by Nucleofection along with Cas9 nuclease and sgRNA against CKAP2. Transfected cells were subsequently treated with Blasticidin for selection of successfully edited cells, and knock-in was confirmed by microscopy visualization of GFP fluorescence. **(b)** Representative time-lapse images of a GFP:CKAP2 knock-in HT1080 cell undergoing a mitotic division. Note that CKAP2 localization in HT1080 cells mirrors that of RPE-1: CKAP2 initially localizes to the centrosomes at the onset of nuclear envelope breakdown (NEBD; red arrows) and is shifted to the chromatin after anaphase onset (green arrows). **(c and d)** Representative immunofluorescence images of RPE-1 cells immunolabelled against CKAP2 (green) and Tubulin (magenta), and stained with DAPI (blue). Note that CKAP2 localization follows a similar pattern in GFP:CKAP2 knock-in cells and immunostained cells. Note also the variability in CKAP2 immunostaining during interphase consistent with a cell cycle dependence of CKAP2 expression (d). In the field of view shown, three interphase cells display CKAP2 immunostaining localized to cytoplasmic microtubules (white arrows), one mitotic cell displays CKAP2 at the spindle (yellow arrow), whereas the remaining interphase cells display no visible CKAP2 immunostaining (d). Time is shown in minutes.

**Supplementary Figure 2.**
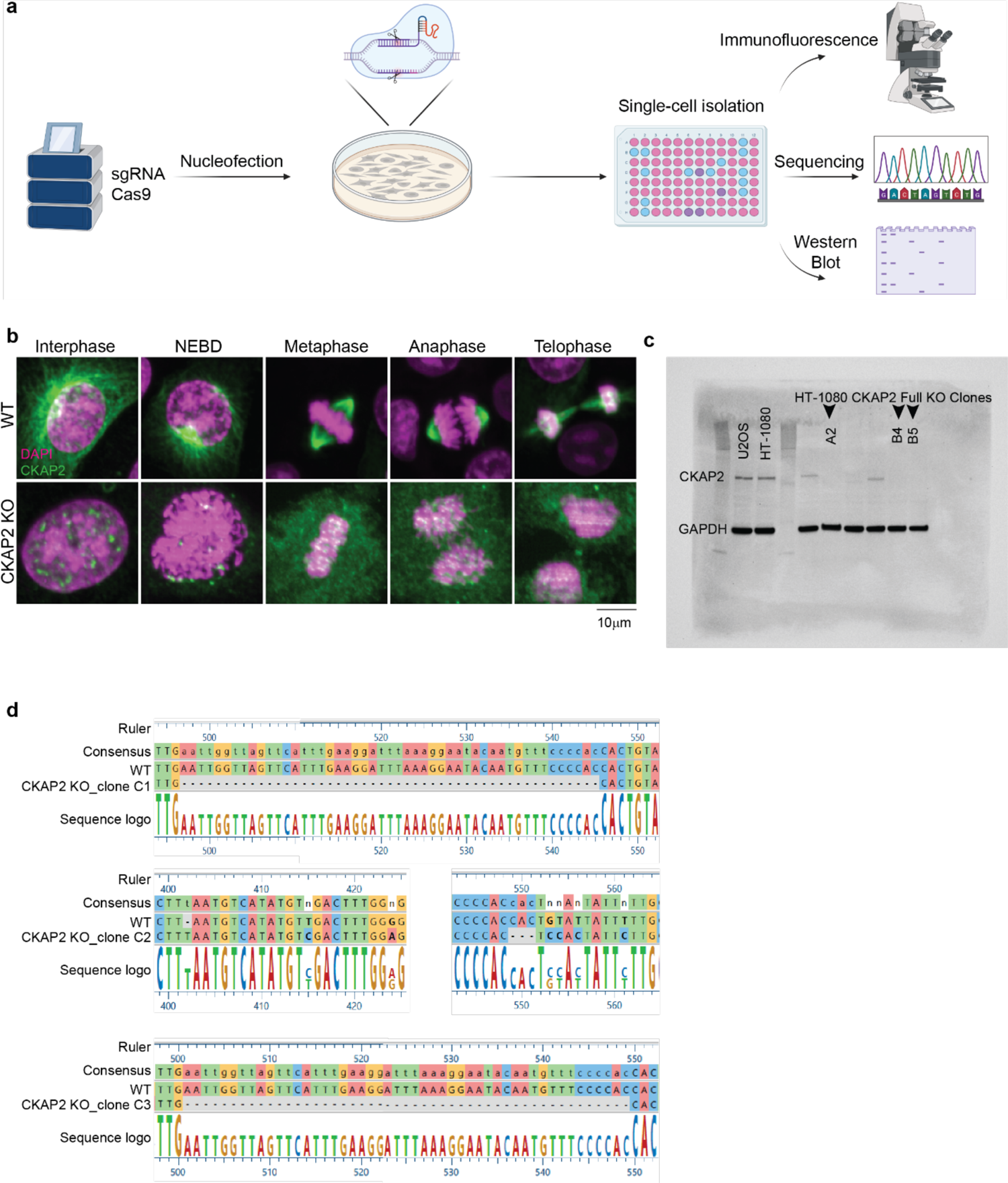
Confirmation of CKAP2 knock-out efficiency. **(a)** Illustration detailing the experimental design for CRISPR-Cas9 knock-out. A Cas9 nuclease and sgRNA against CKAP2 were transfected into either RPE-1 or HT1080 cells by Nucleofection. Clonal isolation was performed into 96-well plates and knock-out efficiency was confirmed in three clones by immunofluorescence, DNA sequencing and Western Blot. **(b)** Representative immunofluorescence images of wild-type (WT) and CKAP2 knock-out (KO) cells from HT1080 clone B4 immunolabelled against CKAP2 (green) and stained with DAPI (magenta). Note the absence of CKAP2 staining in CKAP2 KO cells throughout mitosis. **(c)** Uncropped Western Blot for HT1080 WT and CKAP2 KO clones. Three full CKAP2 knock out clones were selected for study (A2, B4 and B5) and GAPDH used as a loading control. **(d)** Sequencing analyses of RPE-1 CKAP2 KO clones C1, C2 and C3 compared to a WT sequence. Sequence comparisons show indel mutations (black dashes lines) in all three KO clones. NEBD = nuclear envelope breakdown.

**Supplementary Figure 3.**
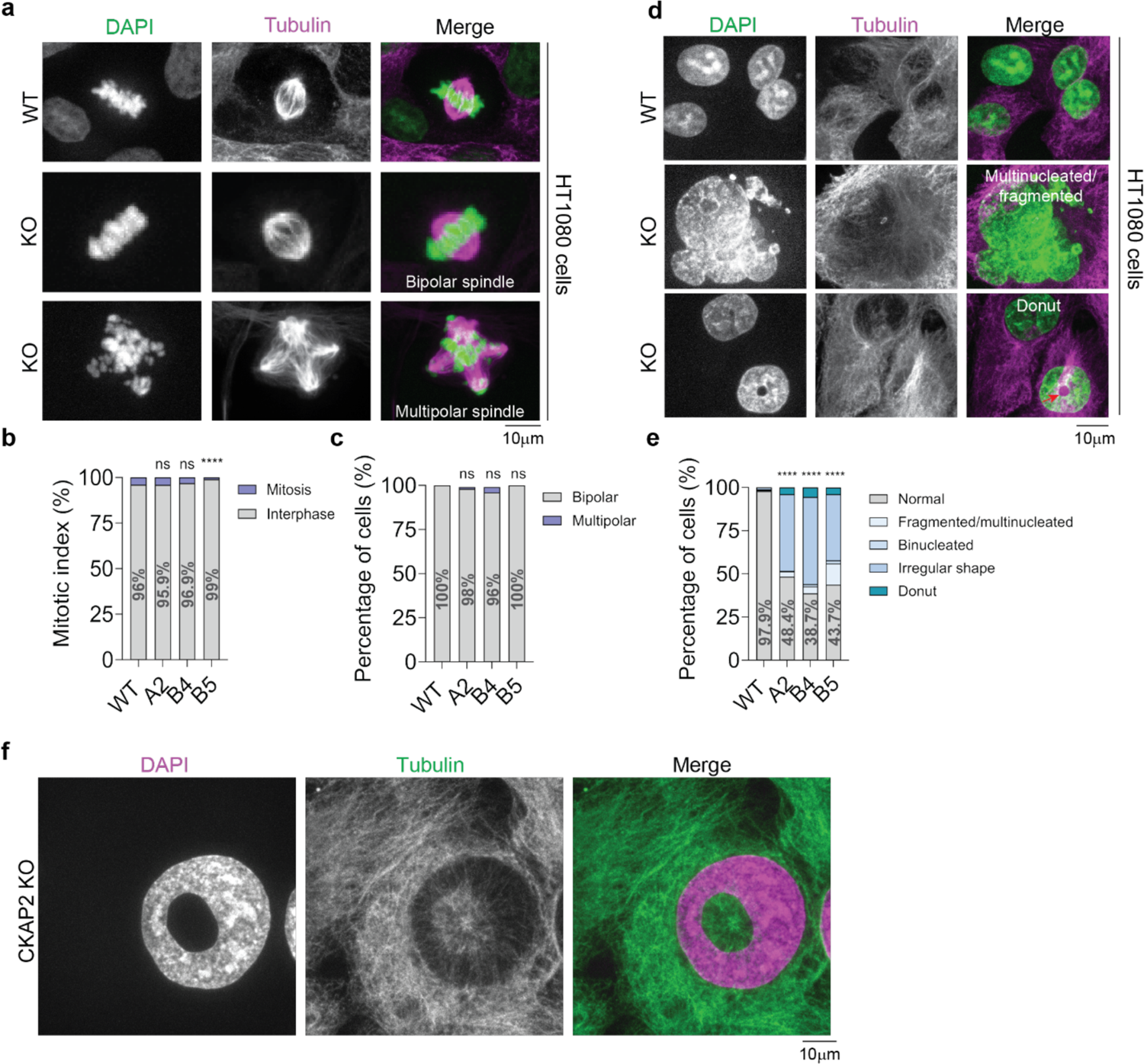
CKAP2 knock-out causes nuclear abnormalities in HT1080 cells. **(a and d)** Representative immunofluorescence images of wild-type (WT) and CKAP2 knock-out (KO) HT1080 cells during mitosis **(a)** and at interphase **(d)**. Note that KO cells display nuclear abnormalities such as multinucleated/fragmented nuclei and donut-shaped nuclei (red arrow in d). **(b)** Quantification of percentage of WT (n = 3800 cells) and CKAP2 KO HT1080 cells (A2 n = 1647 cells; B4 n = 2075 cells; B5 n = 540 cells ****p<0.0001) in interphase vs. mitosis (mitotic index). **(c and e)** Quantification of mitotic spindle **(c)** and interphase nuclei **(e)** abnormalities in HT1080 WT (mitosis n = 151 cells; interphase n = 3649 cells) and KO clones A2 (mitosis n = 66 cells; interphase n = 1581 cells) B4 (mitosis n = 65 cells; 2010 cells) and B5 (mitosis n = 5 cells; interphase n = 535 cells; ****p<0.0001). **(f)** Representative immunofluorescence image of a HT1080 CKAP2 knock-out (KO) cell immunolabelled against Tubulin (green) and stained with DAPI (magenta). Note the presence of a centrosome that appears to be embedded inside the hollow donut surface.

**Supplementary Figure 4.**
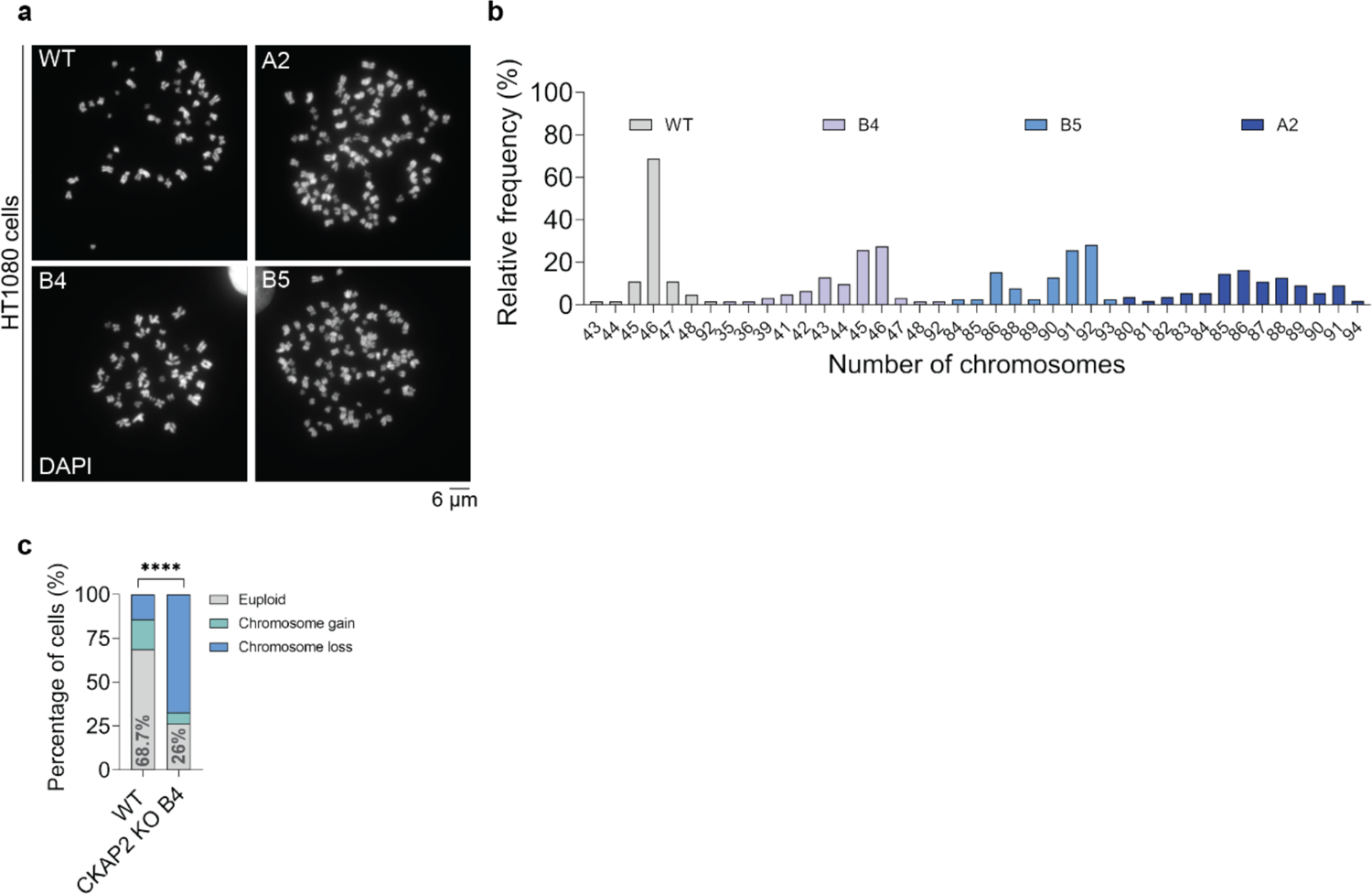
Loss of CKAP2 leads to aneuploidy in HT1080 cells. (a) Representative metaphase spread images of HT1080 WT and KO clones A2, B5 and B4. (b) Frequency distribution of ploidy counts in HT1080 WT (n = 64 spreads) and KO clones A2 (n= 55 spreads), B5 (n = 39 spreads) and B4 (n = 62 spreads). (c) Percentage of HT1080 WT (n= 64 cells) and CKAP2 KO cells (n = 62 cells; ****p<0.0001, clone B4) cells displaying chromosome gains and chromosome losses.

**Supplementary Figure 5.**
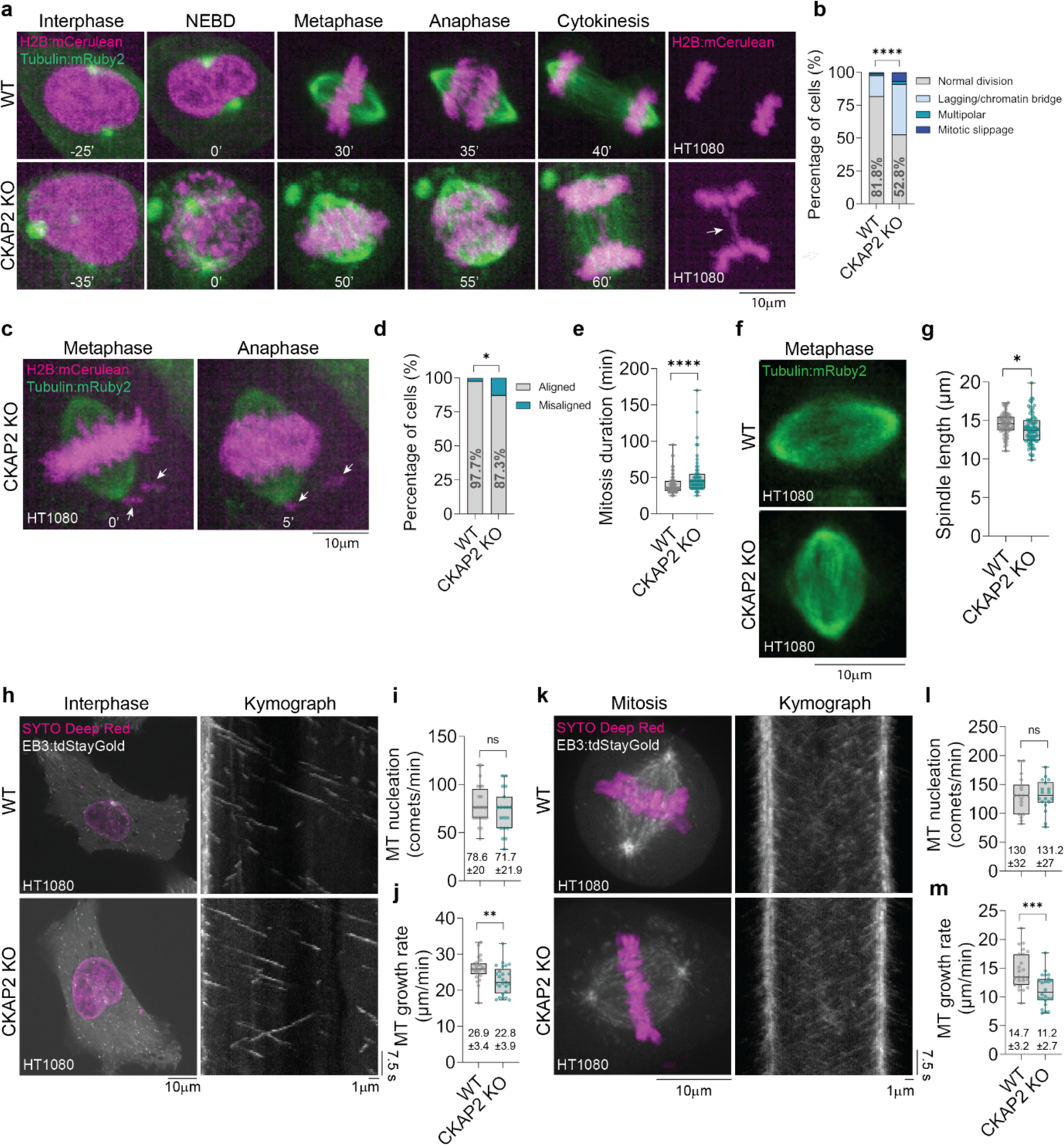
CKAP2 knock-out disrupts chromosome segregation and microtubule dynamics in HT1080 cells. **(a)** Representative time-lapse images of HT1080 wild-type (WT) and CKAP2 knock-out (KO) clone B4 co-expressing H2B:mCerulean (magenta) and Tubulin:mRuby2 (green) and undergoing a mitotic division. Note that the CKAP2 KO cell displays a chromatin bridge (white arrow). **(b)** Quantification of chromosome segregation errors in WT (n = 88 cells) and KO cells (n = 89 cells; p<0.0001). **(c)** Representative time-lapse images of a HT1080 CKAP2 KO cell co-expressing H2B:mCerulean (magenta) and Tubulin:mRuby2 (green) and displaying misaligned chromosomes that remain near the spindle pole at anaphase onset (white arrows). **(d)** Percentage of cells displaying aligned vs. misaligned chromosomes at anaphase onset in WT (n = 86 cells) and KO cells (n = 87 cells; p=0.0178). **(e)** Quantification of mitosis duration as measured by the time from nuclear envelope breakdown (NEBD) to the first frame of anaphase onset in HT1080 WT (39.93 ± 12.7 min n = 76 cells) and KO cells (51.3 ± 25.5 min n = 59 cells; p<0.0001 **(f)** Representative snapshots of metaphase spindles in HT1080 wild-type (WT) and CKAP2 knock-out (KO) cells. **(g)** Quantification of metaphase spindle length measured at the last frame prior to anaphase onset in HT1080 WT (14.56 ± 1.4 µm n = 69 cells) and CKAP2 KO cells (13.85 ± 1.9 µm n = 72 cells; p= 0.0160; clone B4). **(h)** Representative snapshots (left) and kymographs (right) of HT1080 wild-type (WT) and CKAP2 knock-out (KO) interphase cells expressing EB3:tdStayGold (grey). **(i and j)** Quantification of microtubule nucleation from centrosomes **(i)** and growth rates **(j)** in WT (nucleation: n = 24 cells, growth: n = 25 cells) and KO interphase cells (nucleation: n = 19 cells p=0.2911; growth: n = 25 cells p= 0.0039). **(k)** Representative snapshots (left) and kymographs (right) of HT1080 wild-type (WT) and CKAP2 knock-out (KO) mitotic cells expressing EB3:tdStayGold (grey). **(l and m)** Quantification of microtubule nucleation from centrosomes **(l)** and growth rates **(m)** in WT (nucleation: n = 18 cells; growth: n = 23 cells) and KO mitotic cells (nucleation: n = 18 cells p=0.9042; growth: n = 24 cells p=0.0002). Time is shown in minutes. Statistical significance relative to WT group. Ns = non-significant at p ≥ 0.05. Measurements reported as average ± standard deviation.

**Supplementary Figure 6.**
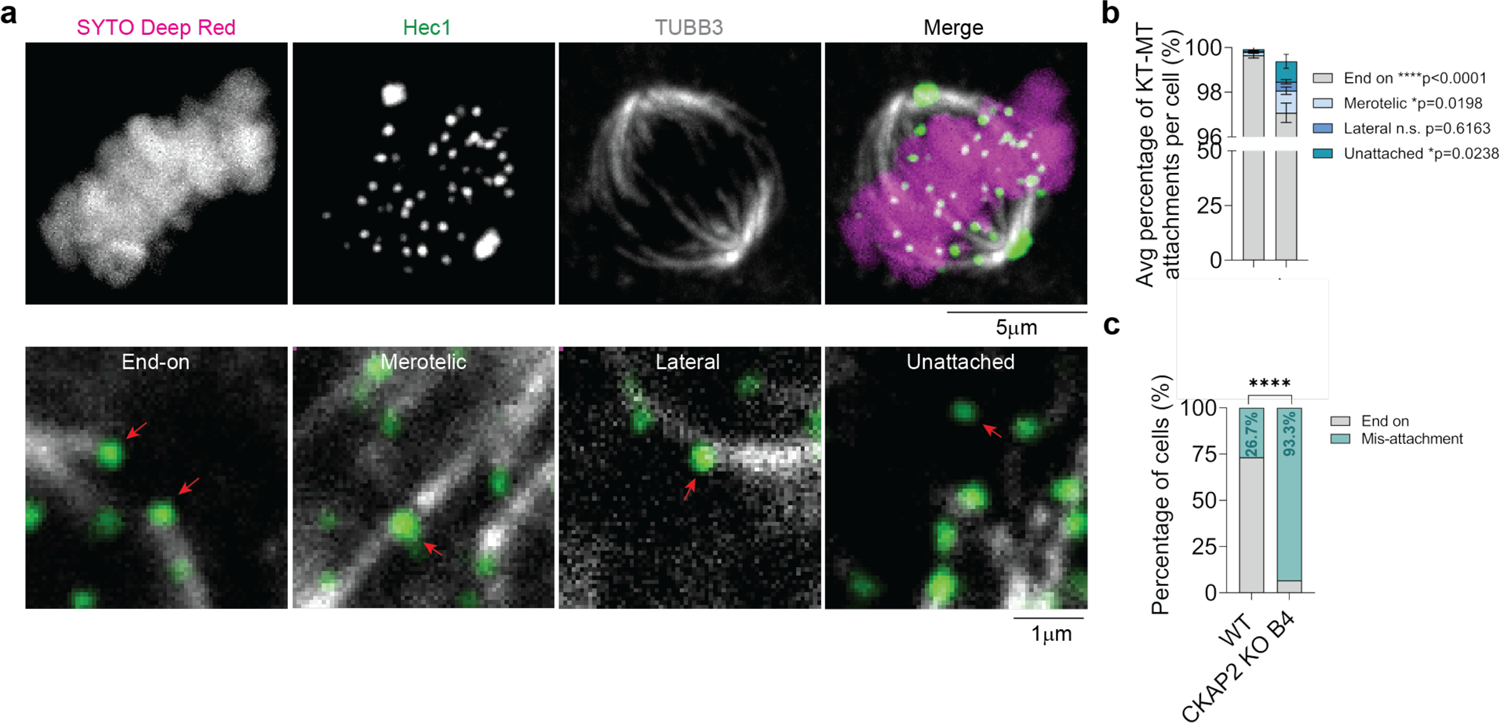
CKAP2 knock-out increases kinetochore-microtubule mis-attachments in HT1080. **(a)** Representative immunofluorescence images of cold-stable kinetochore-microtubule attachments in an HT1080 cell immunolabeled against Hec1 (green) and Tubulin (grey) and stained with SYTO Deep Red (magenta) (top). Zoomed images of end-on, merotelic, lateral and unattached kinetochores (bottom). For merotelic and lateral attachments, only the merotelically/laterally-attached kinetochore are in focus. **(b)** Quantification of average percentage of kinetochore-microtubule attachments per cell in HT1080 wild-type (end-on: 99.65 ± 0.6%; merotelic: 0.14 ± 0.35%; lateral: 0.03 ± 0.1%; unattached: 0.09 ± 0.5% n = 30 cells) and CKAP2 knock-out cells (end-on: 97.1 ± 2.4%; 0.99 ± 0.9%; lateral: 0.4 ± 0.5%; unattached: 0.92 ± 1.7% n = 30 cells). **(c)** Percentage of wild-type (n = 30 cells) and CKAP2 KO cells (n = 30 cells) displaying at least one mis-attached kinetochore (p<0.0001). Measurements reported as average ± standard deviation.

**Supplementary Figure 7.**
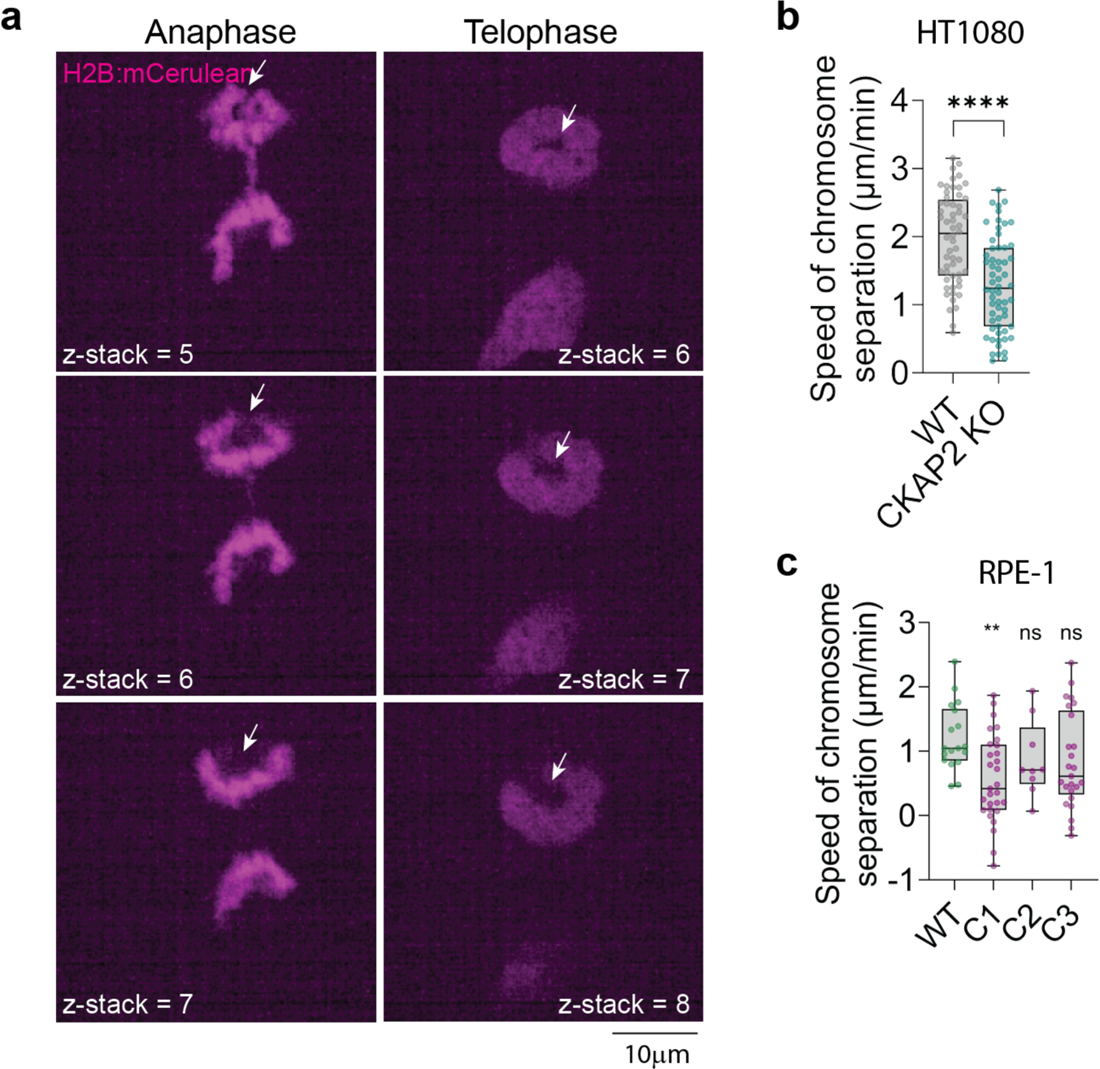
Loss of CKAP2 causes anaphase movement defects. (**a**) Representative time-lapse and z-slices of a HT1080 CKAP2 KO cell (clone B4) displaying abnormal anaphase movement that results in donut nucleus formation. Note that the chromosomes begin to separate but appear to reverse their direction at mid-anaphase, forming a U-shaped chromosome mass followed by donut nucleus formation (white arrows). **(b and c)** Quantification of speed of chromosome separation at anaphase in HT1080 WT (1.97 ± 0.6 µm/min n= 56 cells) and KO cells (1.3 ± 0.7 µm/min n = 60 cells; p<0.0001) **(b)** and in RPE-1 WT (1.2 ± 0.5 µm/min n = 18 cells) and KO clones C1 (0.56 ± 0.6 µm/min n = 29 cells; **p = 0.0096), C2 (0.87 ± 0.6 µm/min n = 9 cells) and C3 (0.84 ± 0.74 µm/min n = 25 cells) **(c)**. Statistical significance relative to WT group. ns = non-significant at p ≥ 0.05. Measurements reported as average ± standard deviation.

